# Climatic refugia in the coldest neotropical hotspot, the Andean páramo

**DOI:** 10.1101/2022.11.21.517325

**Authors:** Gwendolyn Peyre, Catalina Lopez, Maria Daniela Diaz, Jonathan Lenoir

## Abstract

**Aim:** The Andean páramo is the most biodiverse high-mountain region on Earth and past glaciation dynamics during the Quaternary are greatly responsible for its plant diversification. Here, we aim at identifying potential climatic refugia since the Last Glacial Maximum (LGM) in the páramo, according to plant family, biogeographic origin, and life-form.

**Location:** The páramo region in the Northern Andes

**Methods:** We built species distribution models for 664 plant species to generate range maps under current and LGM conditions, using five General Circulation Models (GCMs). For each species and GCM, we identified potential (suitable) and potential active (likely still occupied) refugia where both current and LGM range maps overlap. We stacked and averaged the resulting refugia maps across species and GCMs to generate consensus maps for all species, plant families, biogeographic origins and life-forms. All maps were corrected for potential confounding effect due to species richness.

**Results:** We found refugia to be chiefly located in the southern and central páramos of Ecuador and Peru, especially towards the páramo ecotone with lower-elevation forests. However, we found additional specific patterns according to plant family, biogeographic origin and life-form. For instance, endemics showed refugia concentrated in the northern páramos.

**Main conclusions:** Our findings suggest that large and connected páramo areas, but also the transitional Amotape-Huancabamba zone with the Central Andes, are primordial areas for plant species refugia since the LGM. This study therefore enriches our understanding on páramo evolution and calls for future research on plant responses to future climate change.

## Introduction

The Andean páramo is known as the most biodiverse high mountain region on Earth (Madriñán, Cortés, & Richardson, 2013; Sklenář, Hedberg, & Cleef, 2014; Peyre, 2021). Its extraordinary plant diversification is chiefly due to the uplift of the Northern Andes and its complex history of orographic processes and climatic changes that promoted latitudinal and altitudinal radiations, as well as accelerated *in-situ* speciations, since the Tertiary (Anthelme & Peyre, 2020; Flantua, O’dea, Onstein, Giraldo, & Hooghiemstra, 2019).

In fact, many southern temperate plant taxa, such as *Azorella* and *Oreobolus* (Chacón, Madriñán, Chase, & Bruhl, 2006; Nicolas & Plunkett, 2012) colonized the páramo by taking advantage of the Andean highway migration route that connected the entire cordillera from the Middle Miocene onwards (⁓12 Ma) (Antonelli, Nylander, Persson, & Sanmartín, 2009; Boschman, 2021). In parallel, many tropical lowland plant taxa, such as *Chusquea* and *Elaphoglossum* (Fisher et al., 2009; Rouhan et al., 2004), climbed out of the Amazonian and Chocó cradles and reached higher elevations with the successive phases of mountain-building (Antonelli & Sanmartín, 2011; Hughes & Eastwood, 2006). Moreover, the closure of the Panama Isthmus and the last Northern Andes uplift during the Pliocene (⁓5-2 Ma) led to the increased colonization of the páramo by northern temperate taxa, such as *Festuca* and *Lupinus* (Hughes & Eastwood, 2006; Inda, Segarra-Moragues, Müller, Peterson, & Catalán, 2008).

The páramo then underwent accelerated plant diversification with the glaciation oscillations of the Pleistocene (Flantua et al., 2019; Luebert & Weigend, 2014). During glacial periods, plant species usually migrated downward as the páramo lowered down between 1,000-1,500 m, temperatures decreased, and glaciers expanded (Flantua et al., 2014; Hooghiemstra & Van der Hammen, 2004). During interglacial periods, species migrated upward and resulted isolated as the páramo rose up and formed mountain biogeographic islands. These repeated altitudinal shifts increased speciation rates over short 100 Ka cycles, which in turn contributed to the high plant endemism observed in the páramo today (Londono, Cleef, & Madriñán, 2014; Sklenář et al., 2014).

Understanding how páramo plants responded to past climatic changes is fundamental to improve our knowledge on the biogeography of these unique tropical highlands, but also forecast their vulnerability to future climatic changes (IPCC, 2014; Peyre et al., 2020). To date, an array of methods has been employed to retrieve these ecological responses to past changes in the region.

The common reconstruction approach based on macrofossil and pollen data has been extensively used to describe páramo landscapes and track their floristic composition and physiognomy changes over time (e.g., Flantua & Hoogmiestra, 2018; Hooghiemstra & Van der Hammen, 2004). Moreover, there has been a substantial increase in cladistic and phylogenetic studies over the last decades, shedding new light on the evolution of some important plant groups, for instance, (i) *Lupinus*, Fabaceae (Hughes & Eastwood, 2006; Nevado, Contreras-Ortiz, Hughes, & Filatov, 2018); (ii) *Gentianella* and *Halenia*, Gentianaceae (von Hagen & Kadereit, 2001, 2003); and (iii) *Diplostephium* and *Espeletia*, Asteraceae (Madriñán et al., 2013; Vargas, Ortiz, & Simpson, 2017).

Species distribution models (SDMs) offer a promising and complementary standpoint to other approaches by achieving broad scale biogeographic results for large species pools, while acquitting from potential taxonomic bias and data scarcity issues (Hughes, 2017; Svenning, Fløjgaard, Marske, Nógues-Bravo, & Normand, 2011). In fact, SDMs have frequently being used to hindcast species’ spatial distributions in the past and compare them to present distributions to identify potential climatic refugia where species persisted amidst unfavorable climates (Alvarado & Knowles, 2014; Lenoir et al., 2017; Svenning et al., 2011). Because SDMs can be calibrated with current species distribution data, which is generally easier to gather than paleontological or genetic data, they allow to test broad-scale hypotheses on potential páramo refugia. To date, studies using SDMs to identify refugia have primarily focused on temperate regions (e.g., Rodríguez-Sánchez & Arroyo, 2008; Svenning & Skov, 2004) and more recently on the tropics (e.g., Chan, Brown, & Yoder, 2011; Domic, & Capriles, 2021), but to our knowledge, this is the first attempt to focus on the páramo specifically.

We are interested in understanding how the Pleistocene glaciation cycles, particularly the latest climatic shift between the Last Glacial Maximum (LGM ⁓ 21,000 years ago) and today, created climatic refugia that enabled the long-term persistence of the páramo flora. Given the wide taxonomic spectrum considered, we presumed that the distinctive evolution and ecological range of each plant family played a fundamental part in shaping the potential climatic refugia for its inherent species. Scaling up evolutionary patterns according to biogeographic origin, it is likely that plant species showed coherent spatial patterns between their potential climatic refugia and their migration routes into the páramo. For example, it is probable that strong climatic refugia are to be found at either the south or north ends of the páramo for species of temperate origin, near the páramo-lower forest ecotone for those of tropical origin, and within the region for endemic species.

Moreover, since plant life-forms mirror the functional strategy of species in their environment (Díaz et al., 2016), understanding the distribution of climatic refugia at the life-form level could indicate complementary adaptations and facilitation trends under temporal filtering of the species pool (Bruelheide et al., 2018; Daru, Van der Bank, & Davies, 2018; Mucina, 2019). Consequently, meaningful correlations are expected to be found between climatic refugia for life-forms and macroclimatic features, such as enduring cold environments enabling the long-term persistence of small and resource-saving life-forms (e.g., acaulescent rosettes and cushion and mats) but not tall ones (e.g., upright shrubs). Although spatial trends of potential climatic refugia are undoubtedly diverse among taxa, biogeographic origins, and life-forms, they may nevertheless coincide in specific areas. Accordingly, it is likely that all patterns indicate high values for potential climatic refugia near the páramo ecotone with lower-elevation forests (⁓3,000 m), since this area suffered the least ecological shifts during the Pleistocene glaciation oscillations (i.e., stable páramo free from the temporary occupation of forest and glacier).

In this study, we aimed at identifying potential climatic refugia for plants in the páramo between the LGM and today, by running SDMs for a representative species pool from different plant families, biogeographic origins and life-forms. We modelled species distributions under different SDM settings to draw out both potential (suitable) and potential active (suitable and likely occupied) climatic refugia, and therefore identify key refugium areas for páramo plants in the Northern Andes but also in the strict páramo belt. We forecasted that most plant species would present potential climatic refugia since the LGM. To interpret climatic refugia at the plant family, biogeographic origin and life-form levels, we corrected the resulting refugia maps for potential confounding effects due to species richness. As a result, we were able to draw out the high consensus values for climatic refugia for our species pool, regardless of the current richness spatial pattern in the páramo.

## Methods

### Study area

Our study area encompassed the páramo region in the Northern Andes, covering Venezuela, Colombia, Ecuador and northern Peru (Luteyn, 1999). It consisted of three main zones: i) the northern páramos, down to the Pasto node; ii) the central páramos, down to the Paute-Girón valley and; iii) the southern páramos, i.e., the Amotape-Huancabamba zone, down to the depression of Huancabamba, a known biogeographic barrier for many plant species and relict of the Western Andean Portal (WAP), which had isolated the Northern and Central Andes until the Miocene (Antonelli et al., 2009; Boschman, 2021; Weigend, 2002) (Fig. 1). In terms of elevation, the páramo encompasses all highlands in the region and stretches from the montane treeline to the glacier frontline or mountain top. We assumed that the páramo relative position remained constant over time; but because our research goals involved distinguishing (i) potential refugia in the Northern Andes; from (ii) potential active refugia in the strict páramo region, we defined two páramo spatial extents for the subsequent analyses.

**FIGURE 1:**
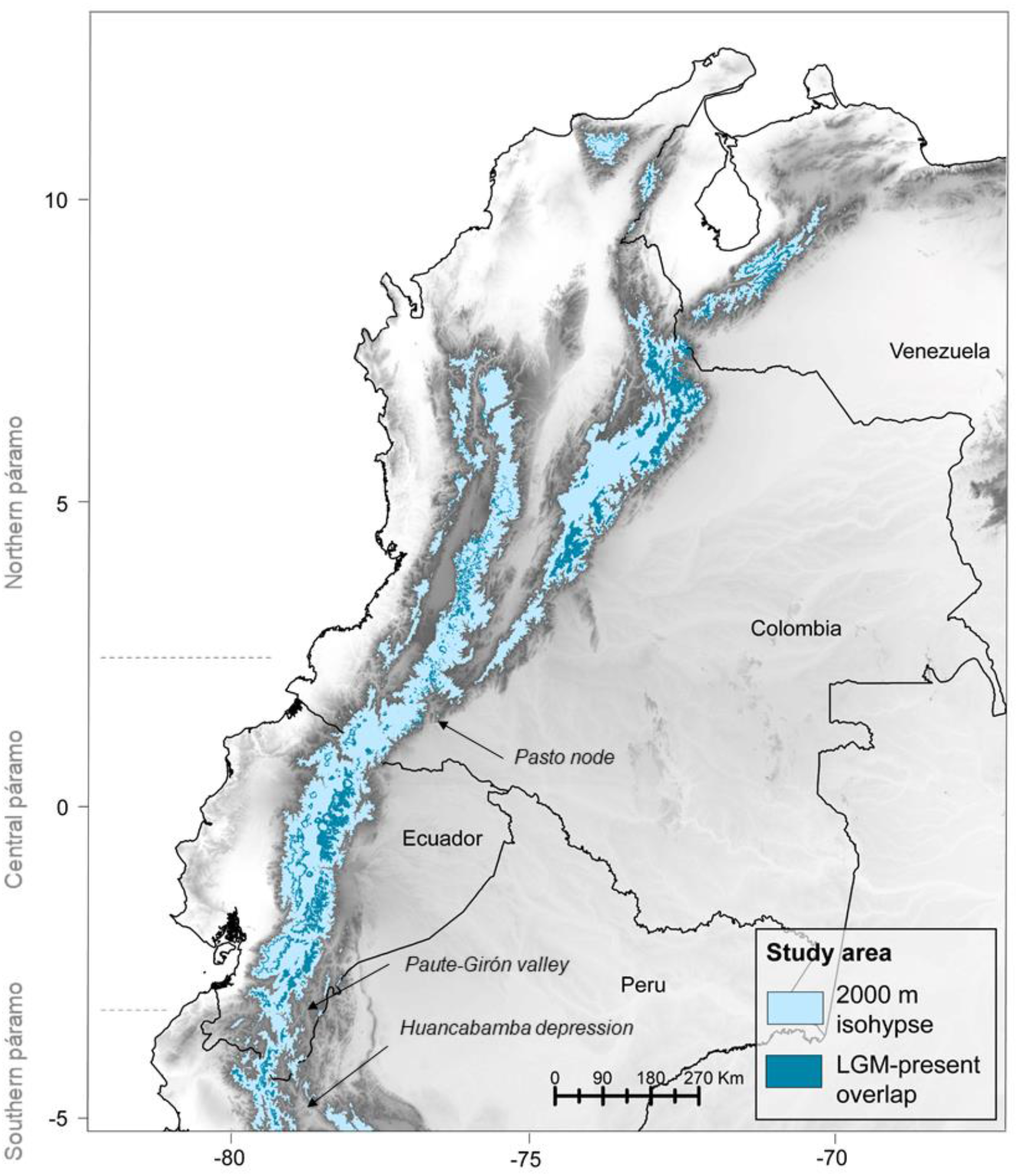
The páramo distribution in the Northern Andes down to the transition with the central Andes (Lat. -5 to -7º). The study area can be distinguished as (i) the Broad Páramo Extent (BPE) delimited by the 2,000 m isohypse (in light blue) during the Last Glacial Maximum (LGM); and the Strict Páramo Extent (SPE), or shared páramo area between the LGM and the present time (in dark blue), delimited by the current Northern Andean treeline and the glacier frontline during the LGM.

The Broad Páramo Extent (BPE), representative of the potential páramo area during the LGM, was set to identify potential climatic refugia for páramo plants in the Northern Andes. The BPE was delimited with the 2,000 m isohypse that embodied the lower páramo elevation limit during the LGM. This frontier was established to account for the 1,000-1,500 m suggested elevation decrease in comparison to today’s páramo treeline, located circa the 3,500 m isohypse (Flantua et al., 2019; Hooghiemstra & Van der Hammen, 2004; Peyre, Osorio, François, & Anthelme, 2021). Setting this lower frontier also prevented our subsequent model predictions to expand and transgress into forest area below. Since we were interested in identifying potential climatic refugia in the Northern Andes, based on climatic suitability only, and regardless of glacier extent, we chose not to constrain the BPE in its upper limit. In contrast, we defined the Strict Páramo Extent (SPE) to assess potential active refugia, hence areas that have likely remained occupied by páramo plants since the LGM. The SPE was delimited to represent the small overlapping surface that has remained constant páramo area between the LGM and today, comprised between the current treeline and the LGM glacier frontline. Glaciers today only cover ∼1% of the páramo region and were therefore deemed insignificant in comparison to the LGM glacier extent (Dussaillant et al., 2019; Peyre et al., 2021). We retrieved the current treeline mask from a recent study on páramo land-covers (Peyre et al. 2021). Establishing the average LGM glacier frontline was challenging because glacier estimates remain scarce and variable throughout the Northern Andes and for this period. Previous studies have set most glacier frontlines between ∼3,000-4,000 m (Angel, Gusman, & Carcaillet, 2017; Mark et al., 2005), but because there was certain multiplicity of methods, timelines and geographic foci (Palacios et al., 2020), we chose to complement this information with a bioclimatic-based approach. We used the 0ºC isotherm, which has been shown to correlate strongly and significantly with the delimitation of glacier fronts in the equatorial Andes (Rabatel et al., 2013), to provide broad-scale standardized estimates of the average glacier frontline throughout our study area during the LGM. To achieve a 0ºC isotherm delimitation of the SPE, we relied on the bioclimatic predictor bio 1 – *Annual mean temperature*, used for the subsequent models and processed for the study area. Bio 1 was prepared for five LGM climate change scenarios based on PIMP3 data (see Section: Bioclimatic data for details). The predictor was first averaged over the different scenarios, then cropped at the 0ºC isotherm, and finally used to fit the SPE (Fig. 1).

### Plant distribution data

We first retrieved the complete set of phytosociological vegetation plots from the VegParamo database (www.vegparamo.com; Peyre et al., 2015), and removed plots located outside the study area as well as plots with inaccurate georeferencing (> 1 km UTM precision). We then edited the dataset by: (i) transforming the phytosociological ordinal scale into presence-absence data; (ii) removing non-vascular taxa; (iii) checking for taxonomic nomenclature using the Plant List (www.plantlist.org); (iv) merging sub-species at the species level; and (v) eliminating nine species that were introduced to the Northern Andes by humans during the past centuries, therefore absent during the LGM. To complement occurrences for species that were underrepresented in the VegParamo dataset (with less than 50 occurrences), we browsed additional data in 18 open-access herbarium databases from Europe and North America for a time-period fitting the VegParamo dataset: 1980-2019. We then checked for occurrence duplicates according to geographic coordinates and data source (based on a 500 m radius from the UTM centroid used in the VegParamo data). Finally, we omitted all species with less than 10 occurrences from the data to reduce overfitting issues in the subsequent models (Merow et al., 2014). The final dataset included 664 species and 3,439 unique observations, 1,986 as VegParamo presence-absence data (shared coordinates among species) and 1,453 as additional occurrences (unique to each species) (Peyre, 2022).

### Species classification

Plant species were classified according to taxonomy (family level), biogeographic origin and life-form. We assigned the biogeographic origin of species at the genus level and following Sklenář, Dušková, & Balslev (2011), complementing data from additional literature whenever necessary. We considered the following categories and subcategories: (i) Tropical – including Neotropical and Wide Tropical; (ii) Temperate – including Holarctic and Austral-Antarctic; (iii) Endemic to the páramo; and (iv) Cosmopolitan – with no clear assignation.

Moreover, we conducted a life-form expert classification based on the 10 categories proposed by Ramsay & Oxley (1997) for páramo grasslands (Fig. 2): (i) Stem rosette; (ii) Basal rosette; (iii) Acaulescent; (iv) Tussock grass; (v) Cushion and mat; (vi) Upright shrub; (vii) Prostrate shrub; (viii) Erect herb; (ix) Prostrate herb; and (x) Trailing herb and liana. Furthermore, we added three extra categories relevant for our study: (i) Epiphyte; (ii) Bamboo, and; (iii) Tree.

**FIGURE 2:**
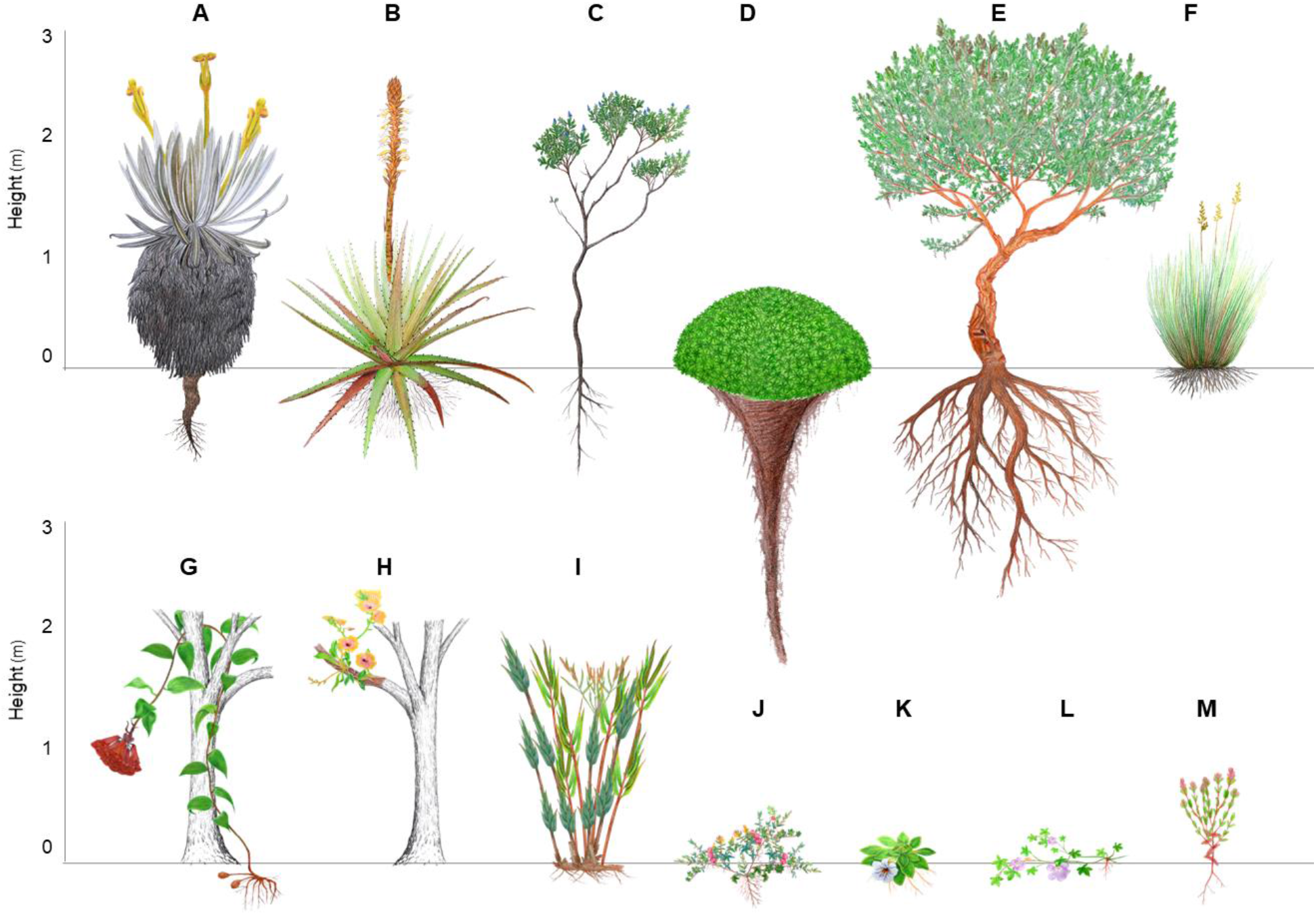
Plant life-forms in the páramo: (A) Stem rosette, illustrated by *Espeletia* (Asteraceae); (B) Basal rosette, illustrated by *Puya* (Bromeliaceae); (C) Upright shrub, illustrated by *Monnina* (Polygalaceae); (D) Cushion and mat, illustrated by *Azorella* (Apiaceae); (E) Tree, illustrated by *Polylepis* (Rosaceae); (F) Tussock grass, illustrated by *Calamagrostis* (Poaceae); (G) Liana and trailing herb, illustrated by *Bomarea* (Alstroemeriaceae); (H) Epiphyte, illustrated by *Telipogon* (Orchidaceae); (I) Bamboo, illustrated by *Chusquea* (Poaceae); (J) Prostrate shrub, illustrated by *Disterigma* (Ericacea); (K) Acaulescent rosette, illustrated by *Viola* (Violaceae); (L) Prostrate herb, illustrated by *Geranium* (Geraniaceae); and (M) Erect herb, illustrated by *Bartsia* (Scrophulariaceae sensu Luteyn et al. 1999, now included in Orobanchaceae).

### Bioclimatic data

Bioclimatic conditions for this study were obtained from the CHELSA project v. 1.1 (http://chelsa-climate.org; Karger et al., 2016, 2017). We downloaded the 19 available bioclimatic variables at a 30 arcsec resolution (⁓ 1 km) for current conditions (1979-2013), and fitted them to our study area, with a first variable set cropped above the 2,000 m isohypse, and a second set delimited with the 3,000 m isohypse. We checked for multicollinearity issues among the second variable set, since it represented closer climatic conditions to the current páramo region. To do so, we used a variance inflation factor (VIF) analysis (*usdm* package; Naimi, Hamm, Groen, Skidmore, & Toxopeus, 2014) and retrieved all variables with a VIF value below 10 (*vif* function). The seven bioclimatic variables selected for both predictor sets were: (i) *bio1* – mean annual temperature; (ii) *bio2* – mean diurnal temperature range; (iii) *bio4* – temperature seasonality; (iv) *bio12* – total annual precipitation; (v) *bio15* – precipitation seasonality; (vi) *bio17* – precipitation of the driest quarter; and (vii) *bio18* – precipitation of the warmest quarter. We downloaded the same variables to represent LGM conditions, relying on PMIP3 data and selecting five general circulation models (GCMs) to encompass sufficient climatic variability (Sanderson, Knutti, & Caldwell, 2015): CNRM-CM5; FGOALS-g2; IPSL-CMA-LR; MPI-ESM-P; and MRI-CGCM3. All past climatic variables were then fitted above the 2,000 m isohypse to represent LGM conditions.

### Species dispersal capacity

Because some of the subsequent models accounted for species’ dispersal capacity, we employed the method implemented in Peyre et al. (2020) to build a dispersal factor.

First, we compiled a basic trait database for the 664 species, building on the available dataset from Peyre et al. (2020). We assessed plant height as the maximum vegetative plant height (in m) at the species level by consulting labeled herbarium data, either in local Andean collections (e.g. ANDES and HNC Herbaria), or online tools, principally JSTOR plants (https://plants.jstor.org) and the Field Museum (https://plantidtools.fieldmuseum.org). Moreover, we retrieved main dispersal mode (zoochory, ballistic, anemochory without flying devices and anemochory using flying devices) and seed mass (in mg) by consulting (i) the Kew Botanical Garden seed collection (http://data.kew.org/sid), and conducting (ii) an extensive complementary literature search on páramo plant seeds (e.g., Frantzen & Bouman, 1989; Melcher, Bouman, & Cleef, 2004). Whenever no data was available for plant height and main dispersal mode at the species level, genus-level information was used. Specific values were averaged over four replicates for all traits, pending data availability (i.e. individual replicates for species-level information, and local species replicates for genus-level information).

Second, we relied on these three significant traits to calculate dispersal kernels emulating the maximum dispersal distance of a species (in m/year) using the *dispeRsal* function (Tamme et al., 2014) in the R statistical software environment (R. 3.5.0). We used the log-transformed value of the maximum dispersal distance, or relative dispersal value λ, to rank the 664 species on a 0-1 scale, from worst (0) to best disperser (1). Finally for each species, λ was projected onto the 3,000 m isohypse-cropped area and used with the *iForce* function to differentiate likely reachable pixels vs. unreachable ones from the current occurrence points (*iSDM* package; Hattab et al., 2017). The final outcome served as species-specific dispersal factor (Peyre et al., 2020).

### Species distribution models

To meet our goals, we performed two series of SDMs, involving distinct projection areas, datapoint filtering and model settings, to emulate species’ potential and realized distributions. We defined the species potential distribution as the area that is environmentally suitable for the species at a given time. In contrast, the species realized distribution represented the fraction of potential distribution that is actually available to the species accounting for its dispersal limitations (Hattab et al., 2017; Peyre et al. 2020; Václavík & Meentemeyer, 2009). The former was later used to retrieve potential climatic refugia, while the latter was used for potential active climatic refugia.

For the potential distribution models, we computed predictions for the 664 species over the BPE area and under current conditions. The models were then projected on the PIMP3 data to reflect LGM conditions (Alvarado & Knowles, 2014; Jackson & Overpeck, 2000). First, in order to best fit the models to the presence/absence data (presences as VegParamo presences and additional occurrences, absences as VegParamo absences), we simply filtered the absence points to remove those whose distance from presence points could dubiously reflect either environmental unsuitability or dispersal limitations only. To do so, we sorted the (i) environmental absences (EA) from the (ii) dispersal-limited absences (DLA) (Hattab et al., 2017), and then restricted our dataset to species occurrences and EA only to better fit the presence data. This was achieved by applying the *pDLA* function, with a α parameter set at 0.5 that considered a 50% probability to classify an absence point as a DLA (*iSDM* package; Hattab et al., 2017). Second for each species, we ran a total of 100 models (25 runs per algorithm) in three steps, model calibration, model validation and ensembling, and model projection. For model calibration, we randomly split the presence/absence data into 75% training and 25% testing datasets (Araújo, Pearson, & Rahbek, 2005). Then, we performed the model on the training dataset using one of the four following algorithms: (i) Generalized Linear Model (GLM); (ii) Random Forest (RF); (iii) Multivariate Adaptive Regression Splines (MARS); and (iv) Artificial Neural Network (ANN) (*biomod2* package; Thuiller, Georges, Engler, & Breiner, 2016). We included four algorithms from different mathematical families to explore the complex relationships between the presence/absence data and bioclimatic information by the means of flexible regression methods, trees and neural networks, and therefore ensure a good-fit of the final ensembled models (e.g., Hallgren, Santana, Low-Choy, Zhao, & Mackey, 2019; Nieto-Lugilde, Maguire, Blois, Williams, & Fitzpatrick, 2018; Norberg et al., 2019). Each model was then validated with the testing dataset. Across the 100 runs, all models whose performance on the true skill statistic (TSS) metric was found ≥ 0.6 were selected for model ensembling (Allouche, Tsoar, & Kadmon, 2006; Marmion, Parviainen, Luoto, Heikkinen, & Thuiller, 2009; Peterson et al., 2011). Following the ensembling process, the model results were projected onto the current predictor set and LGM sets per GCM to obtain the final probabilistic predictions. To ensure comparability between the current and LGM model outputs, we trained the SDMs on the seven bioclimatic variable set cropped at the 3,000 isohypse, but projected the results onto the variable sets cropped at the 2,000 isohypse for both current and LGM conditions. Last, we binarized the predictions using a threshold approach that maximized the sum of the sensitivity and specificity model metrics (*optimal*.*thresholds* function; *PresenceAbsence* package; Freemen & Moisen, 2008;Liu, Berry, Dawson, & Pearson, 2005).

For the realized distribution models, we fitted additional SDMs for current conditions, using the seven bioclimatic predictors and the species-specific dispersal factor computed previously (Svenning et al., 2011). Because macroclimatic variations tend to be subtle in tropical mountain areas, we deemed particularly important to include dispersal into these models to try and best approximate the species’ real distribution (Buytaert, Cuesta-Camacho, & Tobón, 2011; Peyre et al., 2020). The rest of the modelling procedure remained identical to the previous SDMs, except for the: (i) non-filter of absence points between EA and DLA; and (ii) use of the variable sets cropped at the 3,000 m isohypse for both model calibration and projection.

### Final consensus maps

By evaluating the match between a species’ current and past distributions, we drew out its potential and potential active climatic refugia since the LGM. At the species level, we overlapped (i) its current potential and LGM potential binary predictions to produce a potential refugia map (i.e., where climatic conditions have remained suitable for the species in the Northern Andes); as well as (ii) its current realized and LGM potential binary predictions to achieve a preliminary potential active refugia map (i.e., where the species is deemed to have remained since the LGM). These results were first obtained per GCM individually and then averaged across GCMs. Therefore, every species presented two final consensus maps indicating climatic refugia on a 0-1 value scale (i.e., 0 meaning no consensus, and 1 meaning consensus across all GCMs). Then, we stacked the potential refugia maps, already delimited by the BPE, to obtain a consensus potential refugia map for the entire species set, and for species grouped according to plant family, biogeographic origin and life-form. Similarly, we stacked the species’ preliminary potential active refugia maps, but because these results were initially framed by the 3,000 m isohypse, we masked the final stacks with the SPE to emulate consensus potential active refugia solely in the páramo area shared between the LGM and today.

Then, we corrected both consensus maps by the current species richness so to differentiate refugia coinciding with currently diverse areas, hence a potential confounding effect; from (ii) refugia that are independent of richness. First, we produced species richness maps (i.e., for the entire species set and for each species’ group) for current conditions by stacking the probabilistic predictions of the species’ potential distributions (for the potential refugia maps) and species’ realized distributions (for the potential active refugia maps). We relied on probabilistic predictions and not binary ones, since stacking binary predictions tends to overestimate species richness (Dubuis et al., 2011). Second, we rescaled each richness map to the value scale of its corresponding consensus potential and potential active refugia maps. Third, we subtracted the consensus potential and potential active refugia maps with the rescaled richness map and obtained richness-corrected consensus refugia maps. Therefore, positive values reflected on the richness-corrected consensus refugia maps indicated refugia values higher than expected by the current richness (independent refugia), whereas negative values indicated refugia values lower than expected just by the actual richness (coinciding refugia). Finally, we looked into specific responses and identified those species for which the models predicted no consensus values in the study area, namely the no-refugia species.

## Results

Model evaluation proved satisfactory overall, with the RF and GLM algorithms performing best, while ANN fitted the data the least (Table 1). Across the five GCMs and 664 species, models predicted refugia in about 60% of all cases (n = 3,320). Across GCMs, we identified potential refugia for 559 species, corresponding to 84% of the total dataset. The richness-corrected potential refugia map for the 664 species showed consensus for high values in Ecuador and Peru, and to lesser extent in central Colombia, especially towards low elevations (Fig. 3). At the scale of the strict páramo extent, we observed coinciding high values, especially in south-eastern Ecuador and central Colombia.

**TABLE 1.** Model performance according to the Specificity, Sensitivity and True Skill Statistic (TSS) metrics for the four employed algorithms: Artificial Neural Networks (ANN), Generalized Linear Models (GLM), Multivariate Adaptive Regression Splines (MARS) and Random Forest (RF), averaged over their 25 runs for each of the 664 studied species (standard deviation in parenthesis).

|  | **Realized distribution** |  |  | **Potential distribution** |  |  |
| --- | --- | --- | --- | --- | --- | --- |
| Algorithm | Specificity | Sensitivity | TSS | Specificity | Sensitivity | TSS |
| **ANN** | 75.73(14.99) | 78.18(14.70) | 0.67(0.18) | 78.24(11.40) | 78.01 (12.61) | 0.64(0.16) |
| **GLM** | 82.95(13.02) | 89.51(9.23) | 0.77(0.16) | 77.95(14.40) | 87.61(10.59) | 0.68(0.18) |
| **MARS** | 86.46(11.18) | 80.79(13.51) | 0.70(0.17) | 83.82(9.81) | 80.82(11.48) | 0.67(0.14) |
| **RF** | 78.55(18.09) | 82.11(19.63) | 0.75(0.21) | 80.72(14.30) | 80.19(18.91) | 0.71(0.19) |

**FIGURE 3:**
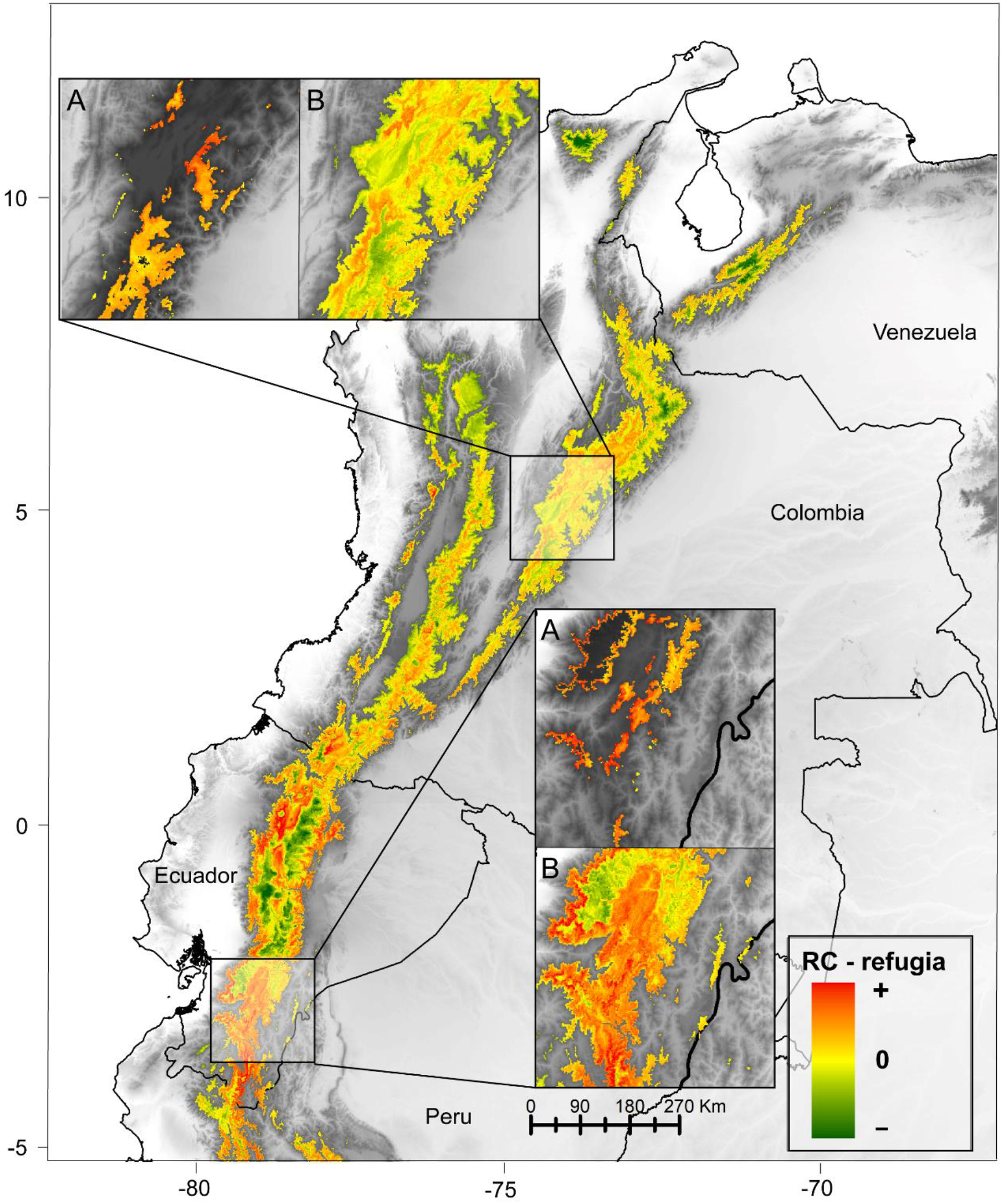
Distribution of consensus potential climatic refugia for páramo plants since the LGM for all species. The map informs on richness-corrected potential refugia (*RC -refugia*), where positive values show refugia independence from actual species richness while negative values show a richness confounding effect. Areas with coinciding potential active refugia are shown in medallions, featuring a zoom in on the exact area from the potential active refugia map (A) and potential refugia map (B) for direct comparison.

Regarding biogeographic origins, we found a slight spatial pattern of increased potential refugia towards the northeastern and southern páramos for temperate species (Fig. 4). However, the highest values for potential active refugia were located in central Colombia and central-eastern Ecuador. Tropical species presented high consensus values for potential refugia throughout the region, especially in southern and northern Ecuador, as well as central Colombia, a pattern that was chiefly repeated on the consensus map for potential active refugia. Finally, high values of potential refugia for endemics were confined to the northern páramos, especially in central Colombia, with the highest values for potential active refugia located in the Colombian eastern cordillera.

**FIGURE 4:**
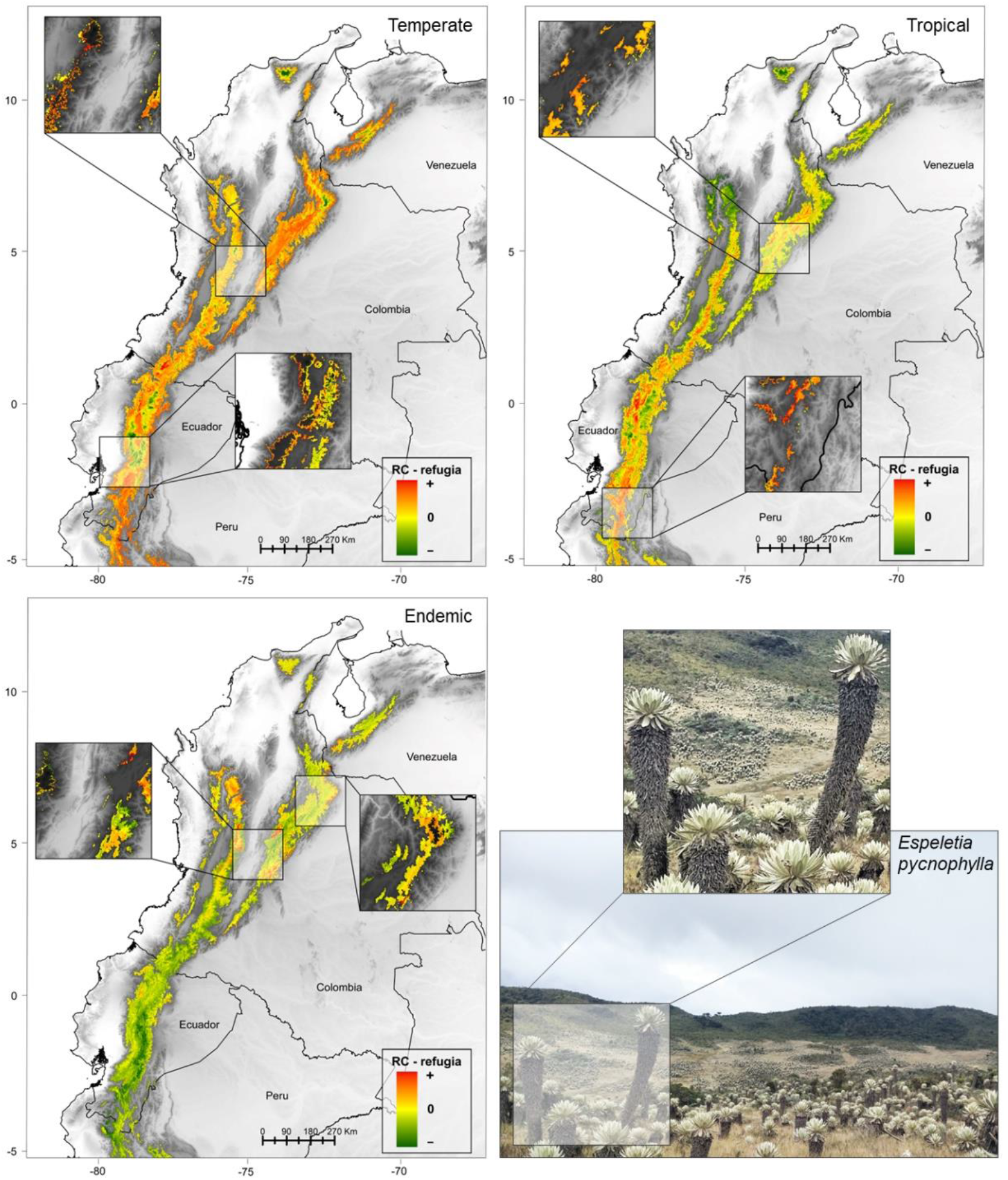
Distribution of consensus potential climatic refugia for páramo plants since the LGM according to the following categories of biogeographic origin: Temperate (Austro-Antarctic and Holarctic), Tropical (Neotropical and Wide Tropical) and Endemic. For each category, the map informs on richness-corrected potential refugia (*RC -refugia*), where positive values show refugia independence from actual species richness while negative values show a richness confounding effect. Areas with coinciding potential active refugia are shown in medallions, featuring a zoom in on the exact area from the potential active refugia map. For the Endemic category, we provided an example of endemic species, *Espeletia pycnophylla* Cuatrec. in the páramo de Paja Blanca (Nariño, Colombia).

Consensus maps for potential refugia at the family level were very contrasted and are presented for the dominant families, Asteraceae, Poaceae, Orchidaceae and Melastomataceae (Luteyn, 1999; Peyre et al., 2015, Fig. 5). Asteraceae potential refugia were mainly located in Ecuador and Peru, a pattern that was repeated for potential active refugia, which were primarily concentrated in the southern páramos. Similarly, we observed important Poaceae potential refugia in Ecuador and Peru, mainly along the eastern slope. However, the main potential active refugia for Poaceae were found in northwestern Ecuador and in the southern páramos. For Orchidaceae, we encountered spread potential refugia over the study area, but potential active refugia stood out in northern Ecuador and central Colombia. There was also a scattered spatial pattern for Melastomataceae, with high values for potential refugia along the western and central Colombian cordilleras and into Ecuador. The consensus map for potential active refugia showed more pronounced values in Ecuador overall.

**FIGURE 5:**
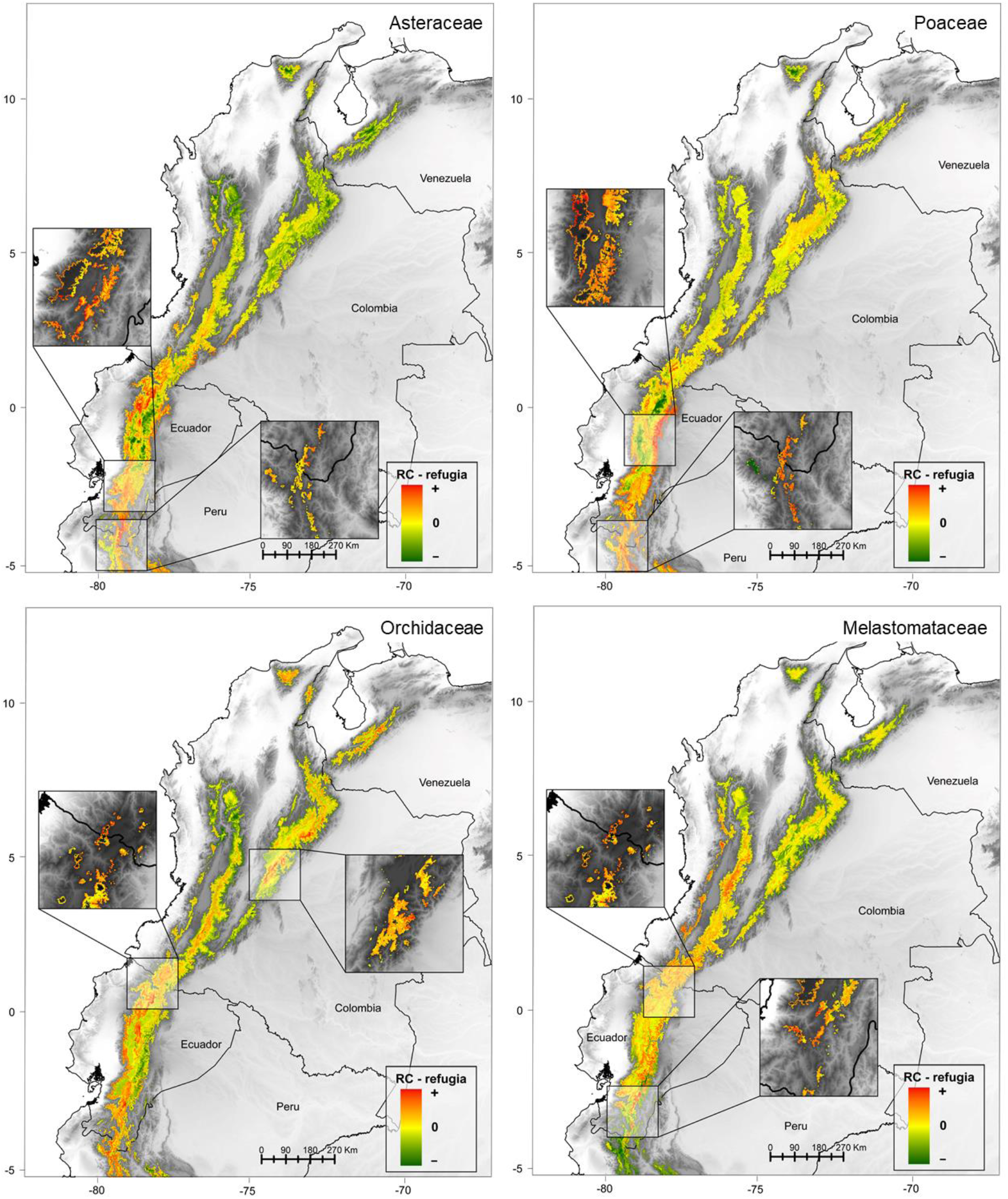
Distribution of consensus potential climatic refugia for páramo plants since the LGM for four dominant plant families: Asteraceae, Poaceae, Orchidaceae and Melastomataceae. For each category, the map informs on richness-corrected potential refugia (*RC -refugia*), where positive values show refugia independence from actual species richness while negative values show a richness confounding effect. Areas with coinciding potential active refugia are shown in medallions, featuring a zoom in on the exact area from the potential active refugia map.

We show the diverse spatial patterns of potential refugia for the most characteristic and abundant life-forms in tropicalpine areas: stem rosette, basal rosette, cushion and mat, and tussock (Fig. 6). For stem rosette species, high values for potential refugia were found at the edges of the study area and concentrated in central Colombia. Regarding potential active refugia, high values were mainly encountered in the same area, but also in Venezuela. For basal rosette species, the highest values for potential refugia were observed in the Colombian eastern cordillera and Venezuela, a pattern that was shared with the map for potential active refugia. Cushion and mat species presented spread high values for potential refugia throughout the region, especially at the edges of its distribution. Nevertheless, we found the highest values for potential active refugia in central Colombia and in central-eastern Ecuador. Finally, we perceived high values for tussock potential refugia in Peru and eastern Ecuador. The tussock potential active refugia were more dispersed, with some coinciding with the potential refugia map in Ecuador, while others differed, as illustrated by the high consensus values encountered in the northeastern páramos.

**FIGURE 6:**
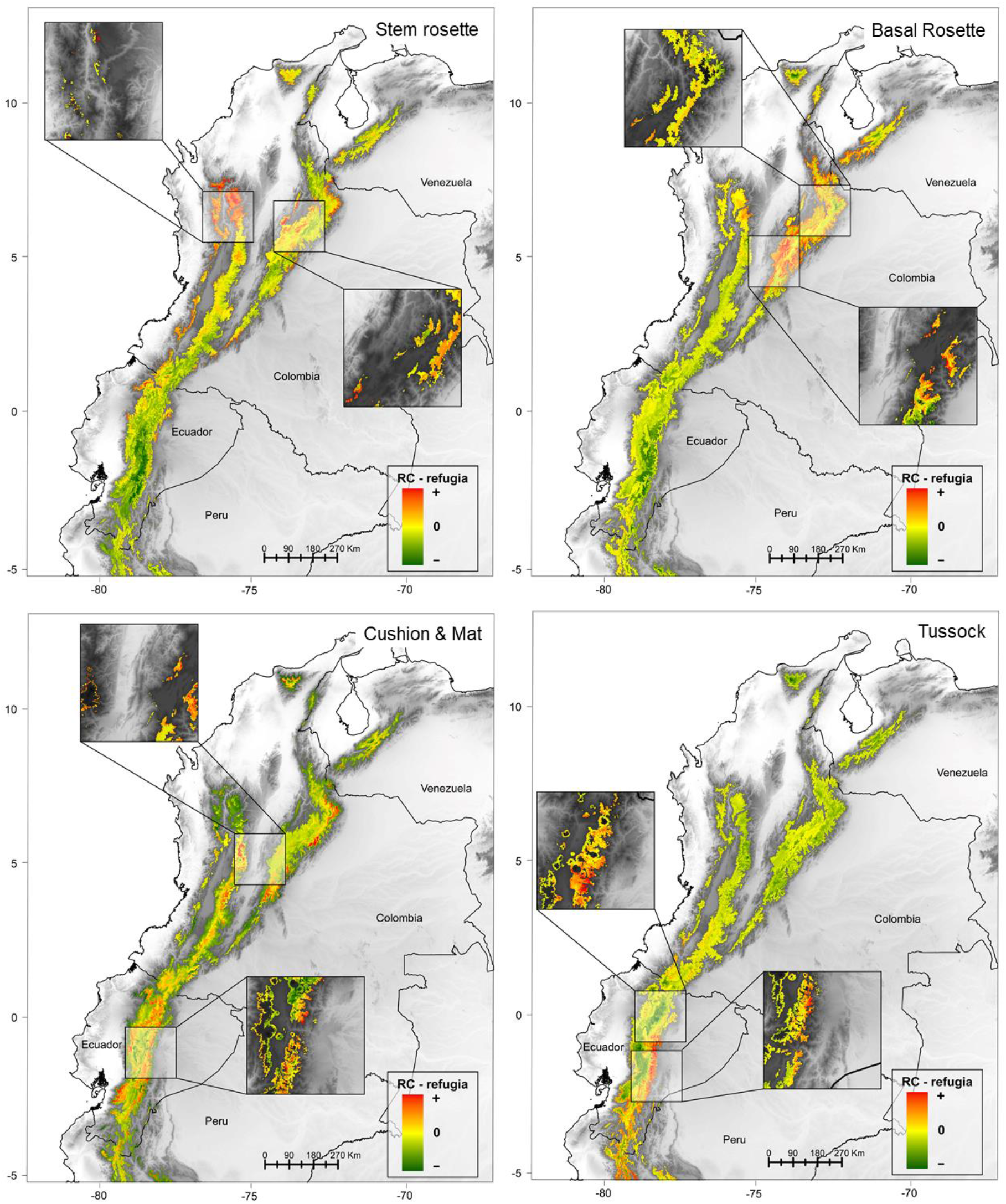
Distribution of consensus potential climatic refugia for páramo plants since the LGM according to four representative and/or dominant life forms: Stem rosette, Basal rosette, Cushion and mat and Tussock. For each category, the map informs on richness-corrected potential refugia (*RC -refugia*), where positive values show refugia independence from actual species richness while negative values show a richness confounding effect. Areas with coinciding potential active refugia are shown in medallions, featuring a zoom in on the exact area from the potential active refugia map.

## Discussion

Repeated periods of habitat isolation and connectivity during the Pleistocene have driven fast diversification in the páramo but also challenged plants’ capacity to persist in a specific area, or refugium (Flantua & Hooghiemstra, 2018; Flantua et al., 2019). Across all studied species, our results suggested that the chances to find potential refugia between the LGM and today were higher in the central and southern páramos of Ecuador and Peru. We observed diverse spatial patterns of potential and potential active refugia according to plant family, biogeographic origin, and life-form, supporting the various evolutionary trajectories that resulted in the Andean lineage diversification (Hughes & Atchison, 2015; Luebert & Weigend, 2014; Madriñán et al., 2013; Pennington et al., 2010).

### Potential climatic refugia in the central páramos

Based on the general consensus maps, we identified the highest amount of potential and potential active climatic refugia in Ecuador, with the edges of the current páramo distribution (both sensu BPE and SPE) being more likely to harbor refugia since the LGM. This finding might be explained by the important surface area and connectivity of the Ecuadorian mountains down to the Paute-Girón valley, which could have acted as a museum of biodiversity during the Pleistocene, through the LGM, and until today, hence helping plant species persist within the páramo despite climatic oscillations (Crisp et al., 2009; Rull, 2011). In addition, the location of the Ecuadorian páramos at mid-distance within the biogeographic region, could have promoted species accumulation from south-north migrations, resulting in important climatic refugia but also local richness (Peyre, Balslev, Font, & Tello, 2019).

According to our results, the chances to find refugia were high in central-eastern and north-western Ecuador, which could indicate easy connections to the humid lowlands, i.e., Amazonia and Chocó. We found partial support for the climatic refugia hypothesis at the life-form level (Daru et al., 2018), due to the selection of emblematic life-forms when displaying our results. Nevertheless, we indeed observed that climatic refugia for resource-saving life forms increased with environmental constraints at high elevation while others decreased. Moreover, Melastomataceae species, most of them upright shrubs, favored the edges of the páramo distribution to find refugia, a pattern that we also observed for tropical species, and which can be explained by the fact that both species groups are strongly correlated to the sub-páramo transition with lower-elevation forests. This is possibly why many species from these categories have a larger proportion of their LGM distribution in the 2,000-3,000 m elevational range than average (60%) in comparison to higher elevations. Indeed, our results supported the initial hypothesis that tropical species could find more refugia along the páramo slopes to facilitate past connections with humid Andean forests, although this pattern did not clearly stand out in the northern páramos (Antonelli & Sanmartín, 2011; Pennington et al., 2010). It is likely that these species escaped the glacial periods of the Pleistocene by remaining close to their original lowland cradles and then performed recent colonization or recolonization of the páramo though enhanced dispersal (Hampe, 2011; Hughes & Eastwood, 2006). In that context, species distribution models that include lower elevations than the 2,000 m isohypse but also account for precise páramo-forest ecotonal dynamics during the LGM might help shed some additional light on these species’ distribution in the past.

Temperate species also presented main climatic refugia in Ecuador and central Colombia, which contradicts our original hypothesis that they would have remained closer to their north and entry points, near Central America and Peru respectively. This finding could relate to complex evolutionary paths of the species considered or early colonization followed by efficient migration throughout the páramo and later distribution shrinkage of these species in Colombia due to the glaciation dynamics. For example, the holarctic *Festuca* species (Poaceae) crossed from northern America before the Panama isthmus formed, probably by island-hopping, and then widely colonized the páramo (Inda et al., 2008; Luebert & Weigend, 2014). Later, *Halenia* and *Gentianella* species (Gentianaceae) likely took advantage of the newly formed land corridor and contributed to the Great American Biotic Interchange between the two continents (Cody, Richardson, Rull, Ellis, & Pennington, 2010). After entering South America and undergoing fast adaptative radiations, their species could have reached and remained in the central páramos (Luebert & Weigend, 2014; von Hagen & Kadereit, 2001, 2003). Moreover, because temperate species are naturally pre-adapted to cold conditions, they are more likely to colonize the páramo than tropical taxa (Sklenář et al., 2011). This theory could help explain why the refugia of tropical taxa were found mostly on the páramo slopes and distribution edges, whereas temperate ones showed more widely distributed refugia.

### Potential climatic refugia in the southern páramos

During the LGM, the southern páramos were already connected and relatively contiguous with the Central Andes (Anthelme & Peyre, 2020; Antonelli et al., 2009). Theory states that the biotic exchange that occurred between the Central and Northern Andes accelerated with the Quaternary glaciation oscillations and promoted plant diversification and endemism within the small and fragmented Amotape-Huancabamba zone (Weigend, 2002). The high density of potential and potential active refugia in this area supports said theory, as illustrated by the páramo-dominant Asteraceae family. Furthermore, low mountain elevations in the Amotape-Huancabamba zone could have exempted or limited glacier occupation (as seen on the SPE, Fig. 1) and promoted species’ persistence since the LGM. This might help explain the high values for both potential and potential active refugia encountered for forest-páramo ecotonal plants. An example would be the northward migration using the Andean highway migration route of austral-antarctic *Calceolaria* species (Scrophulariaceae sensu Luteyn [1999]), which are commonly found in the sub-páramo belt of the southern and central páramos today (Antonelli et al., 2009; Molau, 1988; Sklenář et al., 2011).

### Potential climatic refugia in the northern páramos

The northern páramos of Colombia and Venezuela are rather insular, and their mountains often present small surface areas, which does not clearly appear on the BPE but stands out on the SPE (Fig. 1; Anthelme & Peyre, 2019). The complex topography of the northern páramos has been deemed an important driver of biodiversity changes under strong oscillating climates in the past (Flantua et al., 2019). Potential and potential active refugia for endemics were mostly concentrated in the northern páramos, which could be due to previous island hopping and vicariance events with the glaciation dynamics, using the páramo as species cradle (Antonelli & Sanmartín, 2011; Madriñán et al., 2013). In fact, it is likely that these species took advantage of the relatively stable climate in the inter-Andean valleys (Hooghiemstra & Van der Hammen, 2004; Särkinen et al., 2012) and adjacent humid lowlands to endure glacial periods and then rose again during interglacial periods by performing local recruitment and adaptation (Sklenář et al., 2011). These repeated processes of changing climate and consequent biotic responses presumably promoted the formation of narrow species niches, often resulting in endemism (Madriñán et al., 2013; Pérez-Escobar et al., 2017). An example of rapid diversification for endemics in Colombia and Venezuela is the stem rosette *Espeletiinae* subtribe (Asteraceae), which appeared at the end of the Pliocene and radiated through the Pleistocene in Colombia and Venezuela (Diaz-Granados & Barber, 2017; Valencia, Mesa, León, Madriñán, & Cortés, 2020).

Besides, the northern páramo also harbours certain broad mountain ranges, such as the nucleus parts of the Colombian eastern and central cordilleras, two areas that our results identified as consensus areas for high values of potential and especially potential active refugia. We suspect these areas were able to maintain more plant species over long periods of time thanks to their larger surfaces and higher connectivity in comparison to neighboring ranges (Flantua & Hooghiemstra, 2018). This pattern appeared to have been chiefly driven by several species groups that today thrive in humid páramos, such as orchids. In addition, specific life-forms must have also taken advantage of these areas to prevail under the oscillating climate, for example basal rosettes, e.g. *Puya* and *Eryngium* species, and cushion and mats, e.g. *Azorella* and *Oreobolus* species (Luebert & Weigend, 2014).

### Future perspectives

According to our results, it is possible to identify potential climatic refugia between the LGM and today for most species in our dataset (84%). However, 105 species were predicted with no-refugia. This finding suggests that some of these species might have had non-coinciding but close distributions above the 3,000 isohypse between the two time periods. It is also possible that certain species distributed in the páramo today have migrated from other areas over the last millennia, and most likely risen from their LGM distribution comprised within the 2,000-3,000 m elevational range or even below pending the existence of open vegetation intrusions or forest ecotones. A final hypothesis is that some species persisted locally within páramo microrefugia unidentifiable at the spatial resolution of our analyses (Lenoir, Hattab, & Pierre, 2017; Rull, 2009), as we suspect is the case for the narrowly distributed super-páramo species *Floscaldasia azorelloides* Sklenář & H.Rob. SDMs are useful to locate historic refugia and understand evolutionary dynamics (Alvarado & Knowles, 2014; Svenning et al., 2011), nonetheless, they can overlook specific genetic and population features only available from phylogenies, which we recommend considering in the future (Antonelli & Sanmartín, 2011; Luebert & Weigend, 2014). Future research could also address species distribution coincidences between the LGM, current conditions and predicted future climate change in order to i) identify long-term stability areas in the páramo, as well as ii) scope which species will likely rely most on adaptation vs. migration in the future (Lenoir & Svenning, 2015; Svenning, Eisenhardt, Normand, Ordonez & Sandel, 2015). Not only would these findings contribute significant insight into the páramo evolution and diversity patterns in light of the 6^th^ mass extinction event (Barnosky et al., 2011), but they could also support conservation outcomes.

